# Retrieval of human aversive memories involves reactivation of gamma activity patterns in the hippocampus that originate in the amygdala during encoding

**DOI:** 10.1101/2024.01.18.576178

**Authors:** Manuela Costa, Daniel Pacheco, Antonio Gil-Nagel, Rafael Toledano, Lukas Imbach, Johannes Sarnthein, Bryan A. Strange

## Abstract

Emotional memories require coordinated activity of the amygdala and hippocampus. Human intracranial recordings have shown that formation of aversive memories involves an amygdala theta-hippocampal gamma phase code. Yet, the mechanisms engaged during translation of aversive experiences into memories and subsequent retrieval remain unclear. Directly recording from human amygdala and hippocampus, here we show that hippocampal gamma activity increases for correctly remembered aversive scenes, while exerting unidirectional oscillatory influence within the theta/beta frequency range on the amygdala for previously seen aversive scenes. Crucially, patterns of amygdala high amplitude gamma activity at encoding are reactivated in the hippocampus, but not amygdala, during both aversive encoding and retrieval. Trial-specific hippocampal gamma patterns showing highest representational similarity with amygdala activity at encoding are replayed in the hippocampus during aversive retrieval. This reactivation process occurs against a background of gamma activity that is otherwise decorrelated between encoding and retrieval. Thus, retrieval of aversive memories is hippocampal-centered, with hippocampal activity patterns apparently entrained by the amygdala during encoding.

## Introduction

Episodic memory is typically better for emotional than for neutral events^1^. One central question is whether emotional episodic memory reflects an augmentation of the same neurobiological mechanism(s) underlying neutral memory, or whether a qualitatively different process is engaged. In humans, lesions of the hippocampus impair both neutral and emotional episodic memory, whereas selective lesions of the amygdala leave neutral memory intact and reduce emotional memory performance to the same level as that for neutral stimuli^2^. Functional MRI studies, although lacking the temporal resolution to measure electrophysiological signals, also point to emotional memory enhancement involving co-participation of the amygdala and hippocampus^3,4^.

Using direct intracranial recordings from humans with drug-resistant epilepsy, we recently described coordinated activity between amygdala and hippocampus during aversive, but not neutral, memory encoding^5^. However, successful encoding of both neutral and aversive stimuli was associated with a common manifestation: increased gamma activity in the hippocampus. The additional component to emotional memory encoding was that amygdala theta activity appeared to coordinate hippocampal gamma power to emotional stimuli. Furthermore, successful emotional encoding depended on the amygdala theta phase at which hippocampal gamma peaked; the phase difference, when considered in the time domain, was related to the lag between peaks of amygdala and hippocampal gamma activity. Thus, the contribution of the amygdala to successful emotional encoding includes a modulation of, and temporal coordination of, hippocampal gamma activity. This unique contribution suggests the presence of a qualitatively different process during emotional and neutral episodic memory.

In addition to tracking memory-related activity as a function of encoding success, a further means to probe the contents of memory is to examine responses to previously encoded stimuli in the context of tests of recognition. Typically, responses are compared as a function of performance, including comparisons of correct hits vs. misses (items previously presented that receive “old” vs “new” responses, respectively) or hits vs. correct rejections (“new” responses of previously unpresented foils). These effects can then be compared between emotional and neutral trials. Recently, the contents of memory have been probed using reinstatement analyses^6^, in which the pattern of activity evoked during retrieval is compared to that measured at encoding to test the similarity between these patterns both at the single stimulus and category (e.g., emotional vs neutral) levels. Converging evidence using functional MRI^7,8^, electrophysiological^9,10^ and magnetoencephalography measures^11-13^ have identified encoding-retrieval similarity: neuronal activity patterns present during encoding that are reactivated during retrieval. Conversely, there is growing evidence for putative alterations in memory representations within various brain regions and memory stages ^14-17^. Thus, some questions remain regarding the interpretation of reactivation-related activity, such as whether BOLD signal measuring patterns of population activity could fail to isolate more transient^14-19^ activity related to the content of memory. An additional question is whether reactivation patterns spread into different brain areas during a period of putative consolidation rather than remaining focal^17,20^.

Here, we investigate amygdala and hippocampal oscillatory responses during recognition of aversive scenes, recorded intracranially in patients from whom emotional memory encoding mechanisms were derived^5^. We ask two central questions: whether encoding and retrieval of aversive information engage the same directionality of amygdala-hippocampus coupling^5^ and whether amygdala and hippocampal responses during recognition reflect reactivation of encoding patterns. We observed that the directionality of amygdala-hippocampus coupling is reversed at retrieval compared to encoding. Reinstatement analyses showed decorrelated encoding and retrieval activity in amygdala and hippocampus selectively for emotional stimuli. However, when representational patterns for remembered emotional pictures were derived from phasic peaks of amygdala gamma activity during encoding, we observed reactivation of these patterns solely in the hippocampus, both during encoding and retrieval. Critically, trial-specific patterns of hippocampal activity showing highest representational similarity to that of the amygdala during successful emotional encoding were subsequently replayed in the hippocampus during successful emotional retrieval. Mechanistically, our results suggest that the amygdala entrains hippocampal patterns during encoding. These hippocampal patterns are later replayed in the hippocampus during retrieval of aversive information.

## Results

Twenty-three participants with drug resistant epilepsy undergoing intracranial recordings encoded and retrieved aversive and neutral scenes, as previously reported^5^. In the encoding session, participants viewed 120 scenes (80 neutrals and 40 aversive) for 0.5 s and were asked if the image pertained to an indoor or outdoor scene. At recognition testing 24h later, an equal number of old and new aversive and neutral scenes were shown and participants made recollection (R), familiarity (K for “know”) and new (N) responses (Fig. 1 a). Adopting the same approach as previously reported^4,5^, memory recollection performance was assessed by comparing the rates of R and K responses for emotional versus neutral items. This analysis revealed an interaction between emotion (aversive vs neutral) and memory (R vs. K), (within-subjects two-way ANOVA, F_(1,22)_=4.39, P=0.048). During both encoding and recognition sessions, we simultaneously recorded from the amygdala (*n*=20 patients) and ipsilateral hippocampus (*n*=14 patients) (Methods, Fig. 1 b). Throughout the manuscript, analyses of electrophysiological data during recognition were limited to previously seen scenes, aiming to investigate the reactivation between encoding and retrieval processes. Analyses of amygdala-hippocampal connectivity were performed on the 14 patients with electrodes in both structures.

**Fig. 1.**
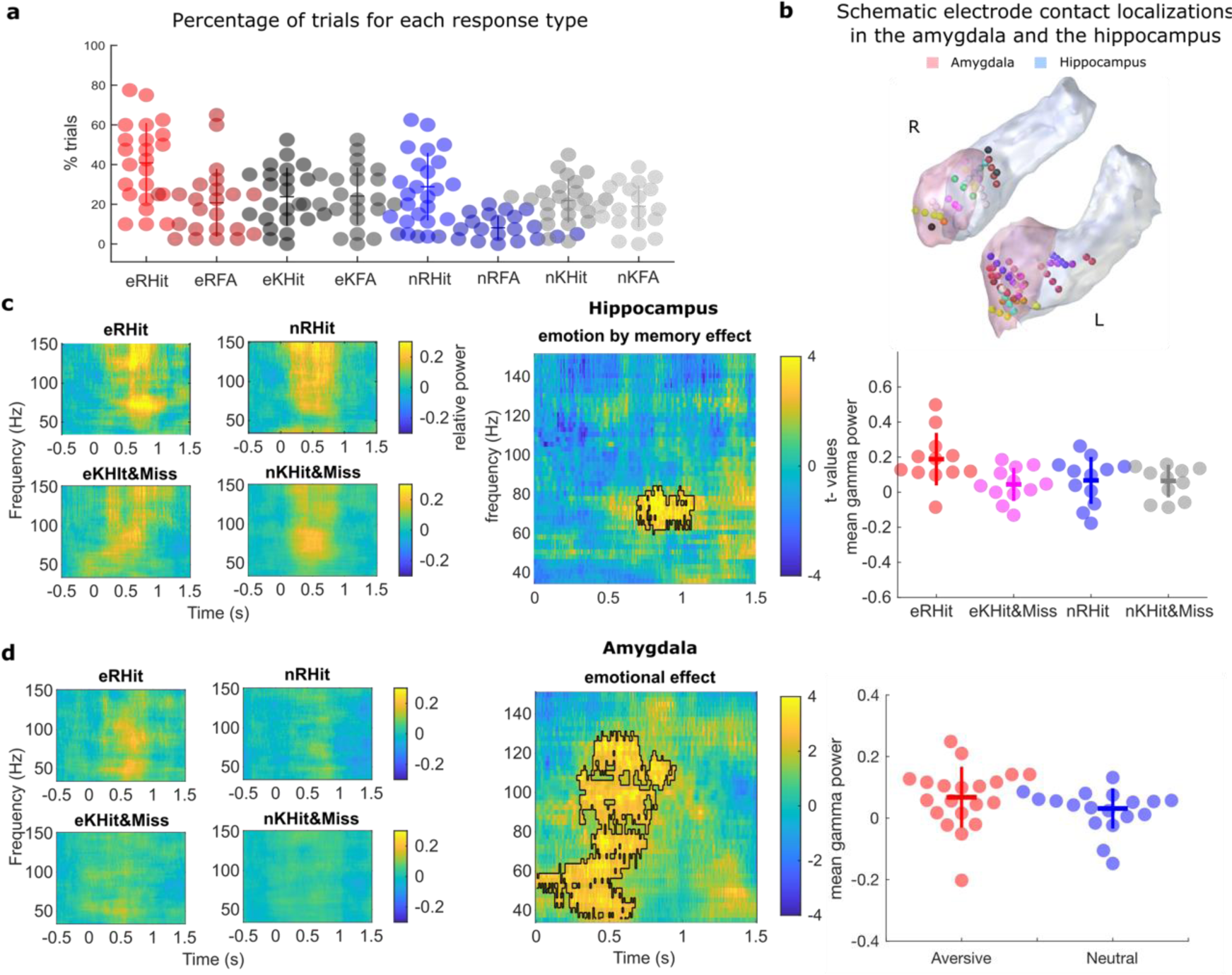
The hippocampus tracks correctly remembered aversive scenes, whereas the amygdala responds to aversive vs. neutral scenes independently of memory performance. **a** Recognition performance (n=23 participants) for aversive (e) and neutral (n) correctly remember (R), remembered false alarms (RFA), correctly known (K) and known false alarms (KFA). Here, and in subsequent plots, each dot represents one patient’s data, and horizontal and vertical line reflects mean and standard error of the mean, respectively. A score of 100% for the eR condition means that there were no false alarms and that every old stimulus was correctly remembered. b Schematic representation of electrode contact localization illustrates the positions of contacts for the 20 participants included in the electrophysiological analysis in the left and right amygdala (pink), along with contacts for the 14 participants in the ipsilateral hippocampus (light blue). Time-frequency plots of hippocampus (c) and amygdala (d) gamma responses (35-150 Hz) for aversive and neutral scenes correctly remembered (eRHit, nRHit) and correct “know” and incorrect “new” responses (eKHit&eMiss, nKHit&nMiss). c Time-frequency resolved test statistics for the comparison of the emotion by memory effect, two-sided paired t-test (cluster-based permutation test). The significant cluster is indicated by a black outline. Scatter plot show the mean hippocampal gamma values in the significant cluster, relative to baseline, for each patient. d same as for c, but for the amygdala time-frequency data. The statistical test shows the comparison aversive vs neutral and the significant cluster (black outline) after applying cluster-based permutation test. Scatter plot shows the mean amygdala gamma power change (relative to baseline) in the significant cluster.

### Retrieval of aversive scenes is hippocampal-centered

We investigated retrieval-related induced responses in the hippocampus and the amygdala by comparing aversive and neutral scenes successfully recollected (eRHit and nRHit, respectively) with scenes that were not. Similarly to previous studies^4,5^, correct “know” (KHit) and incorrect “new” (Miss) responses were collapsed together (eKHit&eMiss, nKHit&nMiss). Aversive scenes correctly remembered (eRHit) elicited higher gamma activity (60-85 Hz) in the hippocampus beginning 0.7 s after stimulus presentation and lasting until 1.1 s (emotion by memory interaction; summed t-value=743.09, P=0.014). Post-hoc t-tests on mean power changes across the significant time-frequency cluster showed a difference for aversive (eRHit vs. eKHit&eMiss t_11_=4.54, P=0.0001, d=1.31, Fig. 2) but not neutral scenes (nRHit vs. nKHit&nMiss t_11_=0.07, P= 0.94, d=0.02). Note that similar hippocampal gamma activity (from 0.7 until 0.9 s, 60-83Hz) was found when comparing the gamma spectral responses to aversive vs. neutral correctly remembered scenes with aversive vs. neutral misses (emotion by memory interaction; summed t-value=475.04, P=0.043, Supplementary Fig. 1a, b), justifying combining ‘KHit’ and ‘Miss’ trials in this and subsequent analyses. At recognition, memory responses can also be assessed by comparing hit and correct rejections. There was a trend towards an interaction when hippocampal gamma activity for aversive vs. neutral correctly remembered scenes was contrasted with those that were correctly rejected (emotion by memory interaction; summed t-value=441.59, P=0.069, Supplementary Fig. 1a-c).

**Fig. 2.**
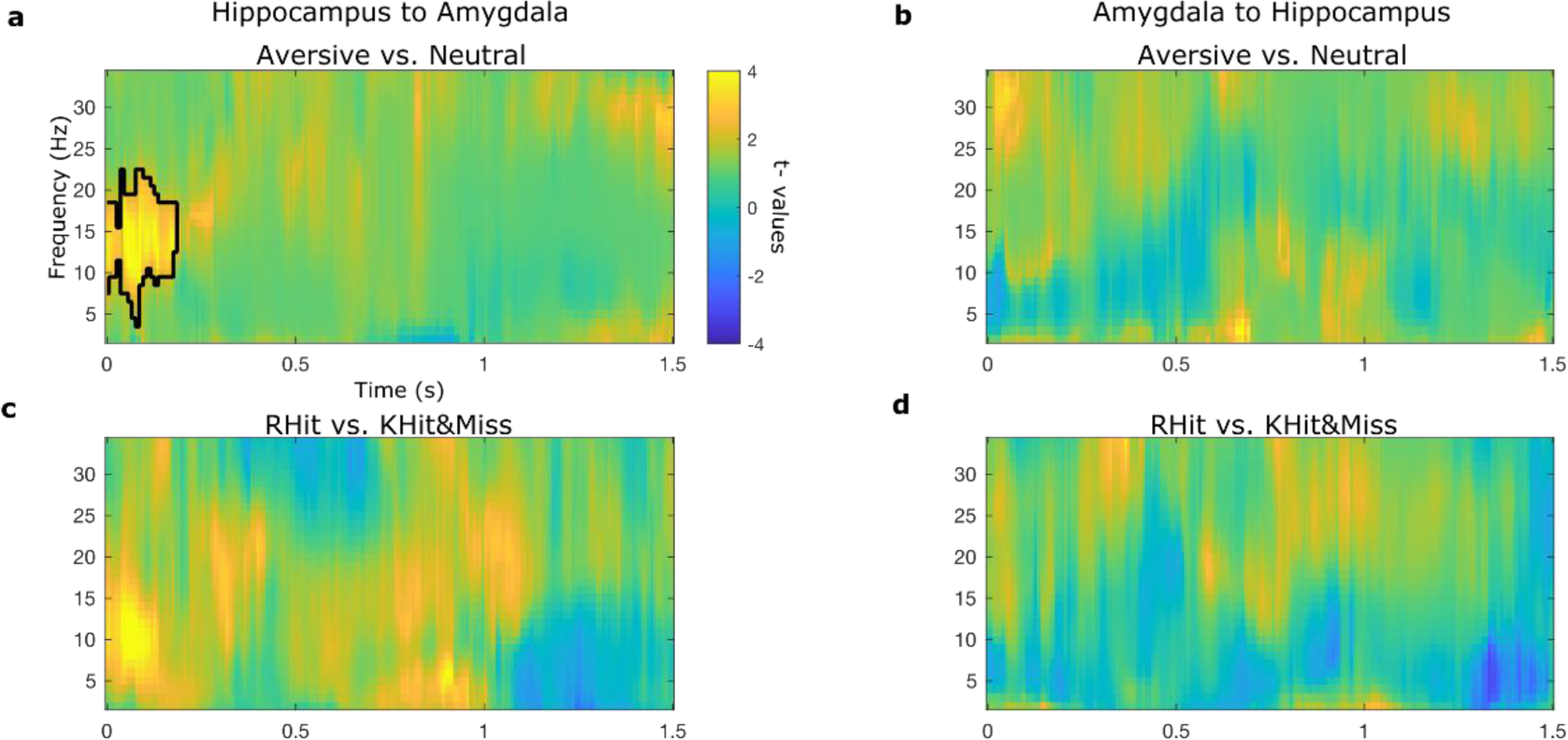
Granger causal influence of hippocampus on amygdala oscillations in the theta to beta range for aversive vs. neutral scenes. **a.** Granger causal influence of hippocampus to the amygdala for previously seen aversive scenes as compared to neutral ones. The significant cluster after applying cluster-based permutation test is outlined in black. **b** same as in **a** but in the direction amygdala to hippocampus. **c-d** Granger causal influence of hippocampus to the amygdala (**c**) and opposite direction (**d**) for RHit vs KHit&Miss comparison.

By contrast to the hippocampus, amygdala gamma activity was not significantly modulated by the recollection of aversive scenes (i.e., an emotion by memory interaction was not observed). A significant increase in gamma power was found while participants observed aversive, but not neutral scenes. This broadband gamma effect (35-130 Hz) started at stimulus onset and lasted until 0.9 s. (main effect of emotion, summed t-value=3887.3, P=0.001, Fig. 1d). Within the lower oscillatory frequencies range (1– 34 Hz), no differences in condition were observed in the hippocampus (Supplementary Fig. 2a), whereas the amygdala showed a significant emotion by memory interaction in the theta range (4 - 6 Hz, from 0.4 until 1.3 s) (summed t-value=493.11, P=0.016). However post-hoc t-tests showed that this was driven by a significant difference only for neutral scenes (Supplementary Fig.2b).

Recollection of aversive scenes therefore elicited different gamma responses in the amygdala and the hippocampus relative to what we observed during encoding^5^. Whereas an emotion by subsequent memory effect is observed in the amygdala during encoding, and emotion by correctly remembered effect is observed in the hippocampus during recognition. We have observed a homologous dissociation with fMRI scanning during encoding and retrieval of emotional vs neutral verbal stimuli^21^.

### Directional influence of hippocampus on amygdala during retrieval of aversive scenes

In rodents, theta oscillations in the amygdala and hippocampus synchronize when fearful memories are recalled^22^. In humans, a stronger directional flow of theta/beta range activity from the amygdala to the hippocampus is observed during encoding when participants viewed aversive scenes compared to neutral ones^5,23^. At retrieval, we first tested the directionality of communication between the amygdala and hippocampus employing frequency-resolved Granger causality for the emotion by memory interaction of previously seen scenes. No significant effects were observed in either direction. However, this analysis revealed higher information flow from the hippocampus to the amygdala for previously seen aversive vs neutral scenes, independent of memory performance. This effect, extending from the theta to the beta band (4-22 Hz) was found from stimulus onset until 0.2 s (summed t-value=655.43, P=0.002), (Fig. 2a). No significant effect was observed in the reverse direction (Fig. 2b). Old scenes correctly recognized as compared to correct “know” and incorrect “new” responses (RHit vs KHit&Miss) did not show a significant effect in either direction (Fig. 2c, d). Thus, Granger causal analysis uncovered higher information flow from the hippocampus to the amygdala specifically for old aversive scenes, directly in opposition to amygdala-to-hippocampal directionality observed during encoding. We note previous evidence showing that, in recognition tasks where novel emotional items were highly similar to those presented during encoding, the correct rejection of these emotional lures involves bidirectional theta oscillations between amygdala and hippocampus^24^.

### Encoding-retrieval global gamma decorrelation selective for emotional memories

There is evidence that memory retrieval involves the reactivation of encoding-related neural activity^7,8,10-13,25,26^. However, an alternative perspective posits that episodic memory operates as a constructive process, entailing neuronal reorganization over time and the transformation of memory content, manifesting as changes in representational patterns between different memory stages^14-17^. Yet, it remains unclear whether amygdala and hippocampal responses observed during recognition reflect reactivation of encoding patterns.

To investigate whether successful emotional retrieval promoted reactivation of encoding activity patterns in amygdala and hippocampus, we employed representational similarity analysis (RSA)^10,27-29^ between the same items presented at encoding and retrieval. Again, we sorted trials into four categories depending on whether participants remembered (R) the aversive or neutral scenes or whether the item received a correct “know”, or incorrect “new” response (KHit&Miss). Specifically, we computed the similarity between encoding and retrieval activity patterns (ERS; Methods, Fig 3a) in the gamma range (between 35 and 150 Hz) in windows of 0.5 s, sliding in 0.4 s (80% overlap). We used Spearman’s rho correlation as our similarity metric and computed the ERS in the amygdala and the hippocampus separately.

**Fig. 3.**
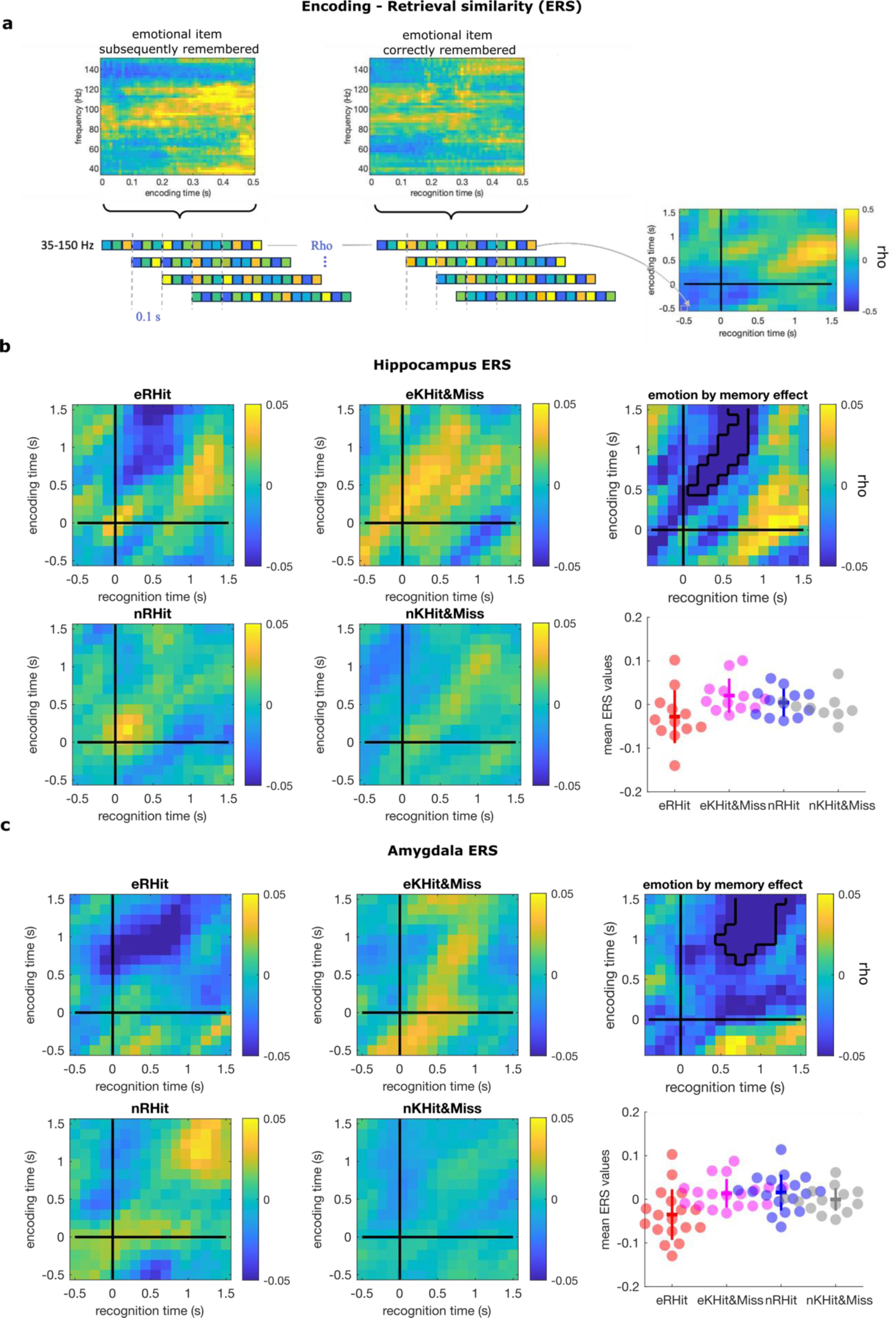
Decorrelated encoding-retrieval gamma (35-150 Hz) activity for aversive scenes in the hippocampus and amygdala. **a** Schematic depiction of the encoding - retrieval similarity (ERS) analysis. Two example epochs of 0.5 for encoding and retrieval are shown. The representational pattern is presented for both sessions as a one-dimension vector which concatenates the frequency (35-150 Hz in 2.5Hz steps) and time dimensions. For each trial, the Spearman correlation was computed between encoding and retrieval from -0.5 to 1.5 s in time windows of 0.5 s, overlapping by 0.4 s, and the correlations averaged within-condition. The encoding-retrieval reactivation map is shown in an example trial, where values in the diagonal represent the reactivation without lag (i.e., the same time period at encoding and retrieval), while off-diagonal values depict lagged correlations. **b** Left: Hippocampus grand average encoding-retrieval reactivation map for aversive (e) and neutral (n) successful encoded and correctly remembered (RHit) for items that received a known/new response (KHit&Miss). Right top: results of the test for the interaction (eRHit-eKHit&Miss) – (nRHit-nKHit&Miss) are shown with the significant cluster outlined in black. The color bar pertains to the correlation coefficient. Right bottom: the mean ERS values for each patient and the four trial types are plotted. Horizontal and vertical line reflects mean and standard error of the mean, respectively. **c** Same as **b** but for the amygdala ERS analysis.

Strikingly, this analysis revealed a negative correlation between gamma activity patterns during encoding and retrieval only for aversive successful encoded and later remembered scenes in both the hippocampus (emotion by memory interaction, summed t-value=-55.73, P=0.048, Fig. 3a) and the amygdala (emotion by memory interaction, summed t-value=-84.34, P=0.017, Fig. 3b). Thus, at least at a macro-level of observation that is induced responses of the local field potential, gamma activity induced at encoding is decorrelated from that induced by the same emotional stimulus at retrieval in both amygdala and hippocampus.

### Encoding-related amygdala patterns around gamma peaks are reactivated only in the hippocampus at encoding and retrieval

Evidence from human recordings suggests that cell assemblies firing at particular frequencies are responsible for encoding episodic events and reinstated during retrieval^30,31^. Gamma bursts have been previously linked to neuronal spiking during memory processes in humans^32^. During aversive memory formation we observed a time-lagged correlation between amygdala and hippocampal gamma peaks ^5^. We thus reasoned that the reactivation process for aversive memory, which was not evident at the level of induced gamma averaged over encoding-retrieval trial pairs, could be better assessed by analyzing neuronal patterns aligned to gamma peaks in the amygdala during encoding. To evaluate this hypothesis, we first identified peaks of high gamma activity (90-150Hz) during encoding in each trial for each condition as previously described^5^. We further assessed reactivation of high gamma activity patterns during encoding (encoding-encoding similarity, EES) and retrieval (encoding-retrieval similarity, ERS) in both the amygdala (Supplementary Fig. 3) and the hippocampus (Fig.4).

**Fig. 4.**
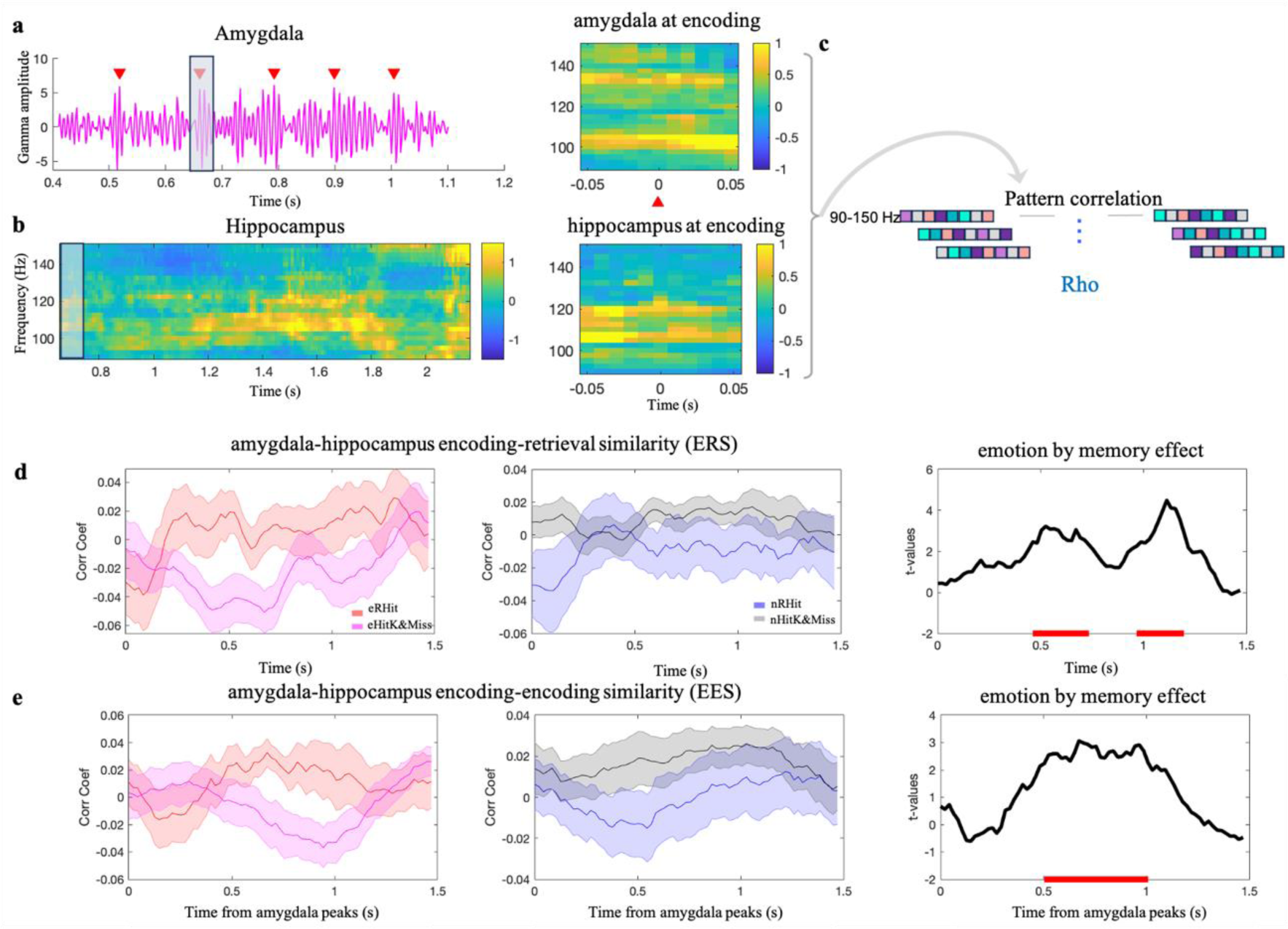
Representational patterns around amygdala gamma peaks at encoding are reactivated in the hippocampus during encoding and retrieval of recollected aversive scenes. Schematic representation of the analysis (**a-c**). **a** Example of one trial showing broadband gamma bandpass filtered between 90-150 Hz. Red arrows point to gamma peaks detected in the amygdala (see Methods). We created a time-resolved activity pattern for each amygdala peak by selecting the activity 0.05 seconds before and after the peak. The opaque vertical rectangle illustrates this selection for one specific peak. One representational pattern from the second peak of the example trial is shown. **b** Hippocampal time-frequency data for the same trial during encoding as in **a**, along with a single representational pattern locked to the time of the amygdala peak. The opaque vertical rectangle represents the hippocampal data windowed 0.05 s before and after the time of the amygdala peak used as an example in **a**. **c**. Each representational pattern was further converted in a one-dimension vector which contain the frequency (90-150 Hz in 2.5Hz steps) and time (0.1 s) resolution. For each trial, in each condition Spearman’s correlation was computed between one pattern in the amygdala and another pattern in the region of interest (amygdala or hippocampus) over time. **d** Encoding-retrieval similarity analysis (ERS): data show the correlation coefficient for eRHit and eKHit&Miss conditions and for nRHit and nKHit&Miss. The x-axis represents the time in the hippocampus from stimulus onset to 1.5 s. T-values are plotted over time. The significant cluster (cluster-based corrected) for the emotion by memory interaction is shown in red. **e** Encoding-Encoding similarity (EES) analysis between amygdala and hippocampus same as for **d**. Here, the x-axis represents time relative to amygdala peaks (0) until 1.5 s after the peak.

Although the current focus is on retrieval of emotional memory, encoding-encoding gamma pattern similarity was also measured, as we reasoned that these results would be helpful in interpreting encoding-retrieval patterns, particularly if these were between-region. Furthermore, there is growing evidence, mainly from human fMRI and scalp EEG studies, that post-encoding awake reactivation strengthens memory^33^. Notably, amygdala representation patterns at encoding were not reactivated in the amygdala at retrieval (Supplementary Fig. 3). By contrast, the ERS analysis revealed a time-period post-stimulus onset from 0.5 to 0.7s and from 0.9 to 1.1s in which amygdala encoding activity patterns were reactivated in the hippocampus only for the aversive scenes correctly recognized (emotion by memory interaction, summed t-value=39.19, P=0.023; second cluster, summed t-value=38.68, P=0.024, Fig 4. d).

The modulation of memory by emotion is generally thought to be a time-dependent process, requiring a period of consolidation^34,35^, although some studies show emotion-induced memory enhancement after only brief delays^36^. The latter observation therefore raises a question as to whether amygdala encoding activity patterns might already be reflected in hippocampal activity patterns during the encoding phase. To address this, we again employed representational similarity analysis, as previously described, but this time to probe encoding-encoding similarity (EES) analysis between patterns of activity in the amygdala and the hippocampus. To define hippocampal patterns at encoding we used the time at which amygdala gamma peaks occurred and again extracted peri-peak activity (±0.05 s) as described for the ERS analysis. We observed patterns of hippocampal gamma activity that mirrored that of amygdala gamma peaks occurring 0.5 s to 1 s earlier, specifically for aversive scenes later remembered (emotion by memory interaction, summed t-value=65.62, P=0.0068, Fig 4. e).

Critically, the reactivation between encoding patterns in the amygdala and the hippocampus at encoding (EES) and retrieval (ERS) were specific to the representational patterns locked to the amygdala peaks, since no significant effects were observed when the signal was locked to random ‘non-peak’ time periods within each individual trial. Results were consistent over 1000 repetitions of random non-peak selection (P-values for the 1000 random selections are reported for the EES analysis in Supplementary Fig. 4a and ERS in Supplementary Fig. 4b).

To further validate the robustness of the observed ERS and EES effects and rule out potential confounds related to the selection of the peaks, we conducted additional control analyses. First, we repeated these tests but increasing the minimum inter peak distance from 0.1 to 0.3 s to account for the autocorrelation of the gamma band activity^37^ in the amygdala and replicated our original results (ERS in the hippocampus: emotion by memory interaction, summed t-value=61.18, P=0.0058; second cluster, summed t-value=54.44, P=0.0092, Supplementary Fig 5a, EES in the hippocampus: emotion by memory interaction, summed t-value=52.65, P=0.012; Supplementary Fig 6a;). Secondly, instead of considering all detected amygdala gamma activity peaks in a trial, we randomly selected one peak per trial. The ERS analysis between amygdala and hippocampus was close to significance (summed t-value=18.38, P=0.075; Supplementary Fig 5b), potentially suggesting that the information carried by more than one peak may be needed for the reactivation process. The EES analysis showed a significant emotion by memory interaction (summed t-value=74.27, P=0.0024; Supplementary Fig 6b).

### Hippocampus patterns at encoding are reactivated during retrieval of aversive scenes

Amygdala high amplitude gamma patterns during encoding reactivated in the hippocampus at encoding and retrieval. A question remained as to whether the hippocampal patterns that mirrored amygdala activity during encoding were also reactivated during the retrieval process. Consequently, we examined reactivation of hippocampal patterns reactivation between encoding and retrieval as a function of the amygdala-hippocampus representation pattern similarity found at encoding. To do so, we used the correlation coefficient score obtained in the encoding-encoding similarity analysis (EES) between amygdala and hippocampus over time. First, for each amygdala pattern derived around amygdala gamma peaks at encoding, we obtained a correlation coefficient reflecting the similarity of this pattern with the hippocampal pattern at encoding over time. Secondly, we determined the time window of maximum similarity between amygdala and hippocampal representational pattern by selecting the time at which the highest correlation coefficient occurred. We next calculated the delay between amygdala gamma peak at encoding and this time point of highest correlation between amygdala and hippocampus and centered hippocampal representation patterns in time to this event (+/-0.05s). Finally, we employed the representational similarity analysis between hippocampal patterns at encoding and hippocampal patterns at retrieval detected from stimulus onset until 1.5 s. A significant emotion by memory interaction was observed (summed t-value=36.16, P=0.021), with the largest difference between conditions identified between 0.7 and 0.9 s after stimulus onset. This finding demonstrates that the hippocampal gamma patterns from the encoding phase that are subsequently replayed during the retrieval of aversive scenes are the patterns present when there is maximal pattern similarity between the amygdala and hippocampus during encoding.

**Fig. 5.**
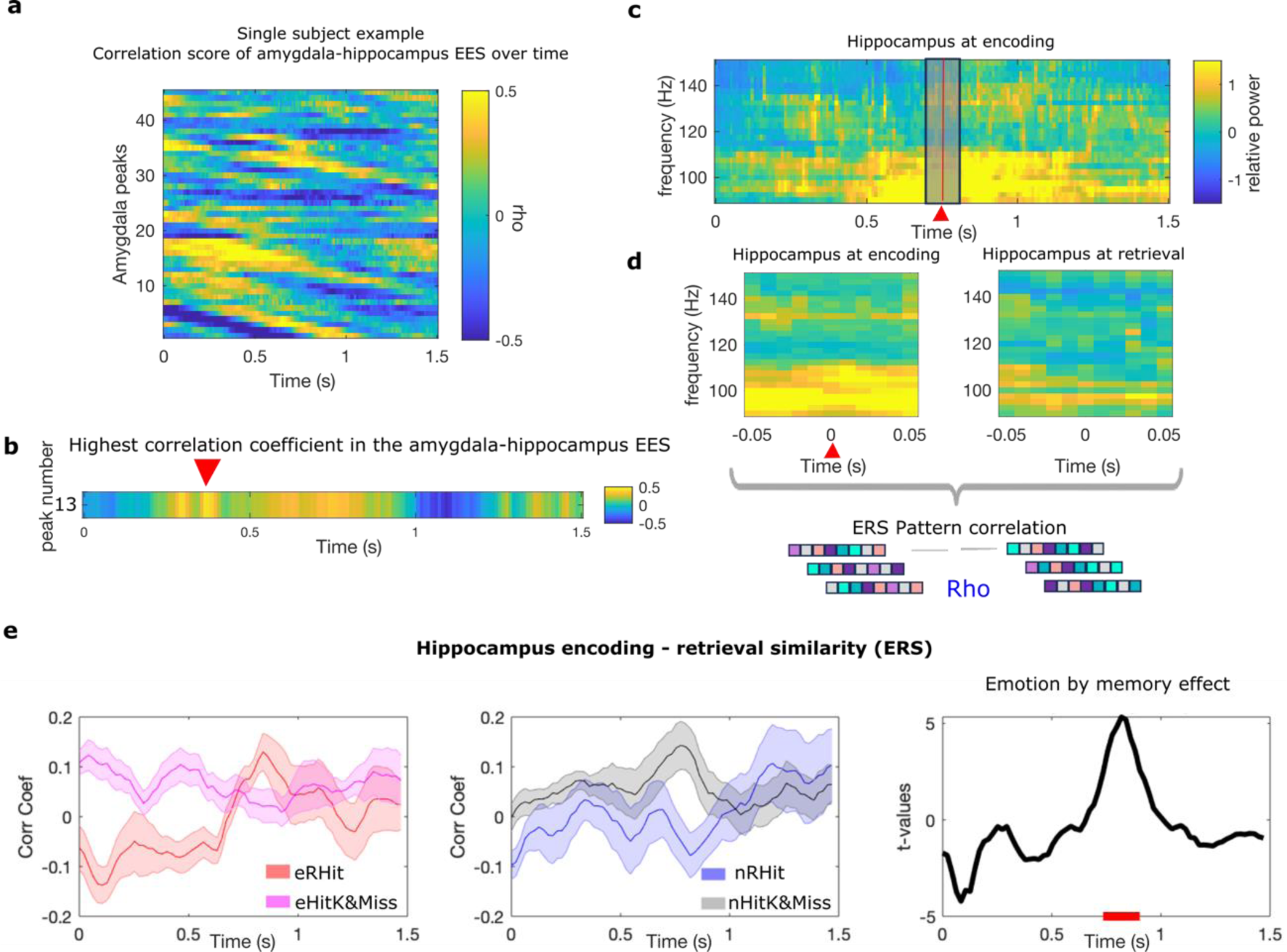
Patterns of hippocampal gamma activity at encoding that are most similar to amygdala emotional encoding activity are reactivated in the hippocampus during emotional retrieval. Schematic representation of the analysis (**a-d**). **a** Example of one subject correlation coefficient of the encoding-encoding similarity (EES) between amygdala and hippocampus over time relative to amygdala gamma peak (x axis). Each row is a peak detected in the amygdala at encoding (y axis). **b** Example of correlation coefficients between amygdala and hippocampus at encoding derived from one amygdala gamma peak. The red triangle points to the highest correlation coefficient. **c** Hippocampal time-frequency data for the same trial where the amygdala peak was detected at encoding. The red vertical line represents the time where the maximum similarity between representational patterns in the two brain structures during encoding was detected (time of the peak + the delay of maximum similarity between the two structures), along with an example of time-frequency data windowed (grey shadow) 0.05s before and after this event **d**. Example of representational patterns included in the RSA analyses. Patterns were converted into a one-dimension vector and the Spearman correlation was computed between one pattern in the hippocampus at encoding and another pattern in the hippocampus at retrieval over time (0.8%-time overlap). **e** Encoding-retrieval similarity analysis (ERS): data show the correlation coefficient for eRHit and eKHit&Miss conditions (left panel) and for nRHit and nKHit&Miss (middle panel). In the right panel, T-values are plotted over time. In all three panels, the x-axis represents the time in the hippocampus at retrieval from stimulus onset (0) to 1.5 s. The significant cluster (cluster-based corrected) for the emotion by memory interaction is shown in red.

## Discussion

Events with emotional significance are generally more memorable compared to neutral ones^38^. However, it is unclear whether fundamentally distinct neurobiological mechanisms are engaged for neutral vs emotional memory formation and retrieval. Both the amygdala and hippocampus are involved in encoding and retrieval of emotional memories^4,5,39^, but currently, only direct and simultaneous recording from both structures can reveal the mechanisms underlying their dynamic interplay in humans. We have previously demonstrated that amygdala oscillations coordinate hippocampal theta and gamma activity during encoding^5^. Here, we provide evidence in favor of qualitatively different amygdala-hippocampal communication underlying aversive memory retrieval.

Through the use of simultaneous intracranial recordings from the human amygdala and hippocampus, we found that during retrieval of previously seen images, hippocampal gamma activity was higher for aversive scenes correctly remembered, whereas the amygdala only showed a general response to aversive vs neutral scenes. The hippocampus unidirectionally influenced the amygdala within the theta/beta frequency range only for aversive scenes as compared to neutral ones. Thus, in contrast to emotional memory encoding, during which the amygdala has been found to direct hippocampal responses ^5^, retrieval of aversive scenes appears hippocampal-centered.

The question of whether memory retrieval involves a reactivation process^8,11^ or rather a transformation or reconstruction of the memory trace^14^ is a subject of ongoing debate^17^. The perspective influenced by the concept of ‘mental time travel’^40^, emphasizes the reactivation of encoded memory representations^10,12,25,26^. By contrast, an increasing number of studies have reported putative transformations in memory representations when examining representational contents in different brain regions between encoding and retrieval ^14-17^. Alternatively, one may argue that different approaches may isolate different aspects of the reactivation process. Neuronal activity could be decorrelated at the macro-level, but the memory content could be better isolated by analyzing neuronal patterns aligned with more transient neuronal activity.

Our first analysis of encoding-retrieval similarity within the gamma frequency range (35-150 Hz) showed that for correctly remembered aversive scenes, both the amygdala and the hippocampus exhibited a decorrelation between encoding and retrieval representational patterns when compared to the rest of trial types (Fig. 3). This finding may reflect the mismatch between gamma frequencies that are employed during successful encoding^5^ (emotion by memory interaction in the amygdala was observed at around 120 Hz) and successful retrieval of aversive scenes (emotion by memory interaction in the hippocampus was observed at around 70 Hz). Thus, this approach did not appear to isolate a reactivation process, instead indicating that– at a global level – gamma activity is different under emotional encoding vs retrieval contexts.

Strikingly, when we restricted the same analysis approach to periods of high amplitude gamma activity, we discovered that these phasic neuronal patterns in the amygdala at encoding did indeed reactivate but only in the hippocampus and only for aversive remembered scenes. These patterns of neuronal activity originating in the amygdala during encoding were replayed in the hippocampus during both the encoding, around 0.4 s after stimulus offset, and retrieval of the same information. The latency of encoding reactivation is consistent with findings from scalp EEG studies, showing a higher degree of reactivation of encoding patterns to pictures later successfully vs unsuccessfully recalled during the post-encoding period, around 0.5 s from stimulus offset^19,41^.

Importantly, the phasic hippocampal gamma patterns that were detected as reactivations of amygdala gamma peaks during encoding, were also reactivated in the hippocampus at retrieval. These results suggest that phasic patterns of amygdala neuronal activity to successfully encoded emotional pictures are mirrored by hippocampal gamma activity (after some lag) during encoding and these encoding-related patterns are reproduced 24h later during the recall of aversive memory^30,42,43^. That is, the reactivation process of phasic amygdala gamma patterns is initiated in the hippocampus during the encoding phase and results in the replay of hippocampal patterns during retrieval in the hippocampus, but not amygdala.

Understanding how re-exposure to aversive stimuli reactivates encoding patterns after 24h is of potential clinical relevance, for example for post-traumatic stress disorder (PTSD), characterized by emotion dysregulation and fear memory ^44,45^, or anxiety disorder. The mechanisms underlying aversive memory reactivation is also relevant to potential therapeutic avenues aimed at rendering these memory traces susceptible to pharmacological^46^ or behavioral interventions. Furthermore, providing a better mechanistic understanding of amygdala-hippocampal communication during encoding, reactivation and retrieval of aversive information – with precise temporal and spectral resolution – can inform the design of novel stimulation protocols that specifically reproduce or disrupt their dynamic communication. These protocols can build on previous evidence that amygdala theta-burst stimulation enhances memory for neutral stimuli^47^, and a recent report that amygdala closed loop neuromodulation reduces symptoms in two PTSD patients^48^.

In summary, our findings provide evidence of a progression of the aversive memory trace over time and shed light on the role of amygdala-hippocampal dynamics that facilitate successful storage and retrieval of aversive memories. This dynamic process is evident in the dissociable responses of the amygdala and hippocampus during memory encoding and retrieval, and opposite directionality of mutual influence during the latter two processes. Critically, our findings demonstrate that amygdala patterns at encoding, detected around gamma peaks, are reactivated in the hippocampus during both the successful encoding of aversive information and its subsequent retrieval. That is, despite a lack of global correlation in gamma activity between the encoding and retrieval of aversive scenes in both the amygdala and the hippocampus, the peak gamma activity patterns originating in the amygdala during encoding are reactivated in the hippocampus during the encoding and retrieval phase. Moreover, hippocampal encoding patterns that mirrored amygdala activity during encoding, at the time of maximum similarity between gamma activity in the two structures, were also reactivated in the hippocampus during emotional memory retrieval process. The overall mechanism emphasizes the prominent role of the amygdala in aversive memory formation. This suggests that the amygdala entrains hippocampal activity during encoding, which, in turn, results in representational patterns that are reactivated in the hippocampus during subsequent retrieval of the same information. These observations have the potential to guide the development of more effective treatments aimed at targeting the amygdala-hippocampal network, ultimately with the goal of enhancing or modifying the content of memory traces.

## Methods

### Participants

In the current dataset 23 patients with refractory focal epilepsy (12 female) participated, with depth electrodes implanted only for diagnostic purposes. All participants had electrodes in the amygdala and 14 also in the ipsilateral hippocampus. 17 participants were recorded at the Hospital Ruber International, Madrid and 6 at the Swiss Epilepsy Center, Zurich, Switzerland. All participants signed informed consent and did not receive financial compensation. The study received full approval from the local ethics committees of the Hospital Ruber Internacional, Madrid, Spain and Kantonale Ethikkommission, Zurich, Switzerland (PB-2016-02055). Three participants were subjected to exclusion from the electrophysiological analysis on the grounds of insufficient signal quality. Results of encoding related responses of 13 of the 23 participants presented here have been previously reported^5^.

### Stereotactic electrode implantation

We conducted a pre-operative contrast-enhanced MRI under stereotactic conditions to map vascular structures before electrode implantation for participants recorded in Madrid at the Hospital Ruber Internacional. This process allowed to calculate stereotactic coordinates for electrode trajectories using the Neuroplan system (Integra Radionics). We used DIXI Medical Microdeep depth electrodes, which are multi-contact, semi-rigid with a diameter of 0.8 mm, contact length of 2 mm, and an inter-contact isolator length of 1.5 mm. For the majority of patients, the implantation followed the stereotactic Leksell method, while three patients underwent implantation using the Medtronic Robot-assisted procedure. For participants recorded in Zurich, we used Ad-Tech depth electrodes with a diameter of 1.3 mm, contact length 1.6 mm, and a spacing of 3 mm between the most medial contact centers. Robot-assisted stereotactic implantation was conducted based on direct targeting in contrast enhanced MRI. Lead localization was verified in post-operative MRI. Depth electrodes were stereotactically implanted into the amygdala, hippocampus, and entorhinal cortex.

### Data acquisition

In Madrid, the intracranial EEG (iEEG) activity was obtained by utilizing an XLTEK EMU128FS amplifier manufactured by XLTEK, located in Oakville, Ontario, Canada. The iEEG data were recorded at a rate of 500Hz at each electrode contact site and they were referenced to linked mastoid electrodes. Intracranial data at the Swiss Epilepsy Center were recorded using the Neuralynx ATLAS system with a sampling rate of 4,096 Hz and an online band-pass filter range of 0.5 Hz to 1000 Hz, all against a common intracranial reference. All data with sampling rate above 500 Hz were down-sampled to 500 Hz.

### Electrode contact localization

To locate the electrodes, we co-registered the post-electrode placement CT scans (post-CT) with the pre-electrode placement T1-weighted magnetic resonance images (pre-MRI) for each patient. To optimize co-registration, we initially skull-stripped both brain images. For CTs, we filtered out voxels with signal intensities between 100 and 1300 HU. For pre-MRI skull stripping, we first normalized the image to MNI space using the New Segment algorithm in SPM8. The resulting inverse normalization parameters were then applied to transform the brain mask into the native space of the pre-MRI. Voxels outside the brain mask with signal values in the highest 15th percentile were filtered out. We then co-registered and re-sliced the skull-stripped pre-MRI to the skull-stripped post-CT. Subsequently, the pre-MRI was affine normalized to the post-CT, effectively transforming the pre-MRI image into native post-CT space. The two images were overlaid, and the post-CT was threshold to visualize only the electrode contacts.

### Electrode contact visualization

To visualize the electrodes using Lead-DBS^49^ (lead-dbs.org), we initiated the process by selecting the electrode model for each patient (Dixi or Ad-Tech). We then performed co-registration of CT scans with MRI using the advanced normalization tools (ANTs), and subsequently, the volumes were normalized into MNI ICBM 2009b nonlinear asymmetrical space based on the preoperative MRI data. Additionally, the software was used to correct for any brain shift that occurred. We initially pre-reconstructed the electrodes using manual reconstruction. The reconstructed electrodes were visually inspected, and if any misalignments were detected, we manually adjusted the reconstruction. This adjustment was based on postoperative CT, ensuring the trajectory was placed as accurately as possible within the center of the electrode artifact. To visualize all the electrodes, we employed the Lead-group software^50^. After this procedure, MNI coordinates were assigned to each electrode contact. In the final step, we plotted only the contacts for each electrode within the regions of interest, either the amygdala or the hippocampus.

### Stimuli

During the encoding session, participants were exposed to a total of 120 pictures—40 with aversive content and 80 with neutral content. These pictures were selected randomly from a larger pool of stimuli, which included 80 high-arousing aversive scenes sourced from the International Affective Picture System (IAPS)^51^ and 160 low-arousing neutral pictures. Aversive scenes involve mutilations, attacks and blood. Of the neutral images, 149 were chosen from the IAPS (depicting household scenes and neutral individuals), while the remaining eleven neutral landscape pictures were sourced from the internet. On a nine-point scale measuring valence, the mean normative IAPS ratings (standard error of the mean) were 5.05 (±0.05) for neutral pictures and 2.04 (±0.05) for aversive pictures. For arousal, the corresponding ratings were 3.29 (±0.06) for neutral pictures and 6.3 (±0.07) for aversive pictures. Given that emotional items tend to be more memorable than neutral ones, the ratio of aversive to neutral stimuli was deliberately set at 1:2 to ensure a balanced number of trials per condition.

### Task

Participants encoded and retrieved after 24h aversive and neutral scenes selected for the International Affective Picture System (IAPS) dataset ^51^. On the second day, they underwent a memory test in which they classified 240 scenes (80 neutral and 40 aversive old, 80 neutral and 40 aversive new) as remembered (R), familiar (K) or new (N). During encoding participants were not informed about the memory test and they were just asked to perform an indoor-outdoor judgment.

### Pre-processing analysis

The FieldTrip toolbox (https://www.fieldtriptoolbox.org)^52^ and Matlab version R2019b (The Mathworks, Natick, MA, USA) were used to analyze intracranial EEG data. By removing signals from nearby electrode contacts in the hippocampus or amygdala, recordings were converted for all participants to a bipolar derivation to optimize local activity^53-55^. If participants had more than one electrode in the amygdala or in the hippocampus, only the one located within the non-epileptogenic zone was used for the analysis. In the case of hippocampal electrodes, we exclusively included the most anterior ones. For each amygdala and hippocampus channel, continuous recording was divided into epochs from -7.5 to 7.5 s with regard to stimulus onset. Trials were inspected visually in the time domain and time-frequency domain to identify artifacts produced by epileptic spikes or electrical noise. When artifacts were detected, the entire trial was discharged.

### Spectral analysis

Each trial’s time-resolved spectrum decomposition was calculated using 7 Slepian multi-tapers for high frequencies from 35 to 150 Hz in 2.5 Hz steps. Slepian tapers were used with windows of 0.4 s width and a 10 Hz frequency smoothing. Low frequencies (<35 Hz) were calculated using single Hann taper, in 5 Hz steps and sliding windows were defined by 7 cycles per frequency step. The relative percentage change with respect to baseline (- 1 to 0.1 s pre-stimulus time) was then calculated. Baseline corrected spectral activity was then averaged over patient trials and channels within the amygdala and the hippocampus.

### Statistical analysis

Trials at retrieval were divided by emotion (aversive or neutral) and response (R, K, N). In line with previous work^4,5^, KN responses were collapsed together. To investigate whether amygdala and hippocampal responses during recognition reflect aversive items reinstatement we restricted the analysis to previously seen scenes only. Two participants were excluded when computing the interaction because only 2 trials in the neutral remembered condition were acquired. We used a cluster-based permutation test^56^ in order to correct the family wise error rate in the context of multiple comparisons across time and frequency dimensions and identify significant interactions or main effects in the time-frequency domain. Using the maximum summation cluster statistic Montecarlo method, we conducted a 2-tailed t test. A paired t-test was performed at each time and each frequency bin for high (35-150 Hz) and low frequencies (0-34 Hz), separately, using a threshold of alpha = 0.025. Clusters were formed by temporal and frequency adjacency at each permutation step, selecting the a-priori time of interest in our data (from 0-1.5 s). Steps in the permutation were performed 10.000 times.

### Time and frequency resolved Granger causality

For the Granger causality analyses, we exclusively considered the most lateral electrode contact pairs (bipolar channels) situated in the amygdala and the hippocampus. This selection was made to ensure a consistent localization of contacts within the lateral regions of the amygdala and the hippocampus across all participants and most probably incorporating into our analyses electrodes positioned within the (baso)lateral amygdala and CA1 region of the hippocampus. The analysis was conducted using the Fieldtrip and BSMART (http://www.brain-smart.org/) toolboxes^57^. The time domain data underwent a low-pass filtering at 85 Hz and was down-sampled to 250 Hz. We calculated time-dependent sets of multivariate autoregressive coefficients in overlapping windows of 0.4 seconds in length, moving forward in 1-time point steps from -0.5 to 1.5 seconds. A model order of 9 which corresponds to a lag of 0.036 seconds was chosen as previously described^5^. Subsequently, Granger causality (2-34 Hz) was computed based on the transfer matrices derived from the autoregressive coefficients. To test statistically the directionality between amygdala and the hippocampus, we compared the Granger coefficients among conditions and use a cluster-based permutation test to quantify the main effect of emotion (aversive vs neutral), the main effect of memory for old items (RHit vs KHit&Miss) and the interaction (eRHit – eKHit&Miss vs nRHit – nKHit&Miss). We performed the test in the direction amygdala to hippocampus and hippocampus to amygdala.

### Encoding-Retrieval similarity (ERS) analysis

We quantified similarity of neuronal patterns in the gamma range (35-150 Hz) when the same item was presented at encoding and retrieval using representational similarity analysis (RSA)^27^. We performed the same analysis in the amygdala and the hippocampus separately. Representational patterns were built concatenating epochs of time- and frequency-resolved power values (47 frequencies from 35-150 Hz in 2.5 steps) in windows of 0.5 s, overlapping by 0.4 s (80%)^10,28,29^. A metric of similarity or reactivation was computed using Spearmańs correlation between encoding and retrieval from -0.5 to 1.5 s. Power data were baseline corrected from -1 to 0.1 s and if more than one bipolar channel was present, the average between channels was performed. For each trial, we defined a window of 0.5 s and included the time course of 35 to 150 Hz power (2.5 Hz steps). Our representational pattern thus consisted of a 2-dimensional matrix containing 47 (frequencies) and 50 (time points) values. To perform the correlations, we concatenated this matrix into a one-dimensional feature vector (of length 47*50=2350). To assess the temporal stability of this pattern we performed correlations across all possible combinations of time points within a trial. This resulted in an encoding x retrieval reactivation map for each trial (Fig. 3a), where the diagonal showed the reinstatement without lag between the two experimental phases and off-diagonal lagged correlations. We performed this map for each trial type (eRHit, eKHit&Miss, nRHit, nKHit&Miss) and tested for main effects and interactions using paired t-tests. We employed cluster-based permutation statistics to correct for multiple comparisons across time points^56^. A paired t-test for each planned comparison was performed at each time point, clusters were formed by temporal adjacency with cluster threshold of alpha = 0.05 and a significant threshold of alpha = 0.025. Permutation steps were repeated 10000 times.

### Patterns locked to amygdala gamma peaks Encoding-Retrieval similarity (ERS) and Encoding-Encoding similarity (EES) analysis

Similar to the above analysis, we used RSA to quantify reactivation of neuronal activity but now selecting patterns of activity around amygdala high gamma peaks (90-150 Hz) during encoding. We selected the high gamma range to avoid an overlap with the effect that we observed in the hippocampus at recognition (55-83 Hz). The time series data was band-pass filtered between 90-150 Hz using a two-pass finite impulse response (FIR) filter with a filter order of 3 cycles of the lower bound. Amygdala gamma peaks were detected in the filter signal from 0.4 to 1.1 s. This is the window in which we observed a significant gamma effect for successful encoding of aversive scenes^5^. Only peaks with a minimum inter peak distance (IPD) of 0.1s were considered. Each peak time was then used as a time-lock to determine frequency-resolved pattern of activity before and after the peak (±0.05 s) in the amygdala during encoding. Both for the amygdala and the hippocampus, we used time-frequency baseline corrected data (see Method above) and only the most lateral amygdala and hippocampus channels in each patient were selected as previously reported for connectivity analyses between these structures^5^. For each frequency resolved pattern, the reactivation measure was calculated using Spearman correlation. We computed similarity analysis between 1) amygdala at encoding and amygdala at retrieval; 2) amygdala at encoding and hippocampus at retrieval (ERS) and 3) amygdala and hippocampus at encoding (EES). To perform the similarity comparisons, we concatenated the representational pattern (25 [frequencies of gamma from 90-150 in 2.5 Hz steps] *11 [0.1s, +/- 0.05s from the peak]) in a one-dimensional vector in window. This was done in a temporally resolved manner proceeding in post-stimulus time steps of 0.1 s (25 *11 = 275) and with an 80% overlap. In our EES analysis we locked hippocampal data to the time at which amygdala peaks occurred. In the ERS analysis, we did not lock the hippocampal activity to the amygdala peaks as the neurophysiological mechanisms at encoding and retrieval may differ in time. Single subject data were then averaged within each condition. Grand averages for each condition were calculated and we tested for the interaction (aversive remember (eRHit) – aversive known/forgotten (eKHit&Miss)) vs. (neutral remember (nRHit) – neutral known/forgotten (nKHit&Miss) applying a cluster-based permutation test^56^.

To test whether the results were selective to the activity occurring only around amygdala peaks, we run the analysis using data not locked to the amygdala peaks. In each trial, we selected random segments of data of the same window length (0.1s) and number as in the original analysis, ensuring that these segments were not locked to any identified peak and had a minim distance of 0.03s from the peaks. We repeated this procedure 1000 times and assessed ERS and EES following the same procedure described in the original peak-locked analysis. Histogram of p-values for each random selection is reported in Supplementary Fig. 4.

To validate our findings, we also conducted a series of control analyses. Firstly, we repeated the test by increasing the minimum inter peak distance from 0.1 to 0.3 s to account for peaks autocorrelation, as reported in previous studies^37^ (ERS: Supplementary Fig. 5a, EES: Supplementary Fig. 6a). Then, we randomly selected only one peak in each trial and repeated the analysis (ERS: Supplementary Fig 5b, EES: Supplementary Fig. 6b). All together, these control analyses allowed us to further validate the specificity of the observed effects and rule out any potential confounding factors related to peak-related data selection.

### Hippocampus Encoding – Retrieval similarity: patterns at encoding were centered around the highest correlation coefficient of amygdala-hippocampus Encoding-Encoding similarity (EES)

We used RSA to quantify reactivation of neuronal activity in the hippocampus between encoding and retrieval. To achieve this, we focused on the moment of greater similarity between representational patterns in the amygdala and the hippocampus. Initially, we computed a correlation coefficient score over time for each amygdala pattern derived around the amygdala peaks at encoding. Then, for each observation we identified the time of greater similarity between the amygdala and hippocampal representational patterns by selecting the highest correlation coefficient. The delay between the timing of the amygdala gamma peak at encoding and the moment of the highest correlation score between the amygdala and hippocampus was then calculated. We constructed hippocampal representational patterns at encoding by considering activity at +/-0.05s from this event. As for the rest of ERS analysis, to perform the similarity comparisons we concatenated the representational patterns (25 [frequencies of gamma from 90-150 in 2.5 Hz steps] *11 [0.1s, +/- 0.05s from the peak]) in a one-dimensional vector. Lastly, we computed the representational similarity between hippocampal patterns at encoding and hippocampal patterns at retrieval in a temporally resolved manner that proceeded from stimulus onset (0) until 1.5 s time steps of 0.1 s (25 *11 = 275) and with an 80% overlap. A metric of similarity or reactivation was computed using Spearman correlation. Single subject data were then averaged within each condition. Grand averages for each condition were calculated and we tested for the interaction (aversive remember (eRHit) – aversive known/forgotten (eKHit&Miss)) vs. (neutral remember (nRHit) – neutral known/forgotten (nKHit&Miss) applying a cluster-based permutation test^56^.

## Data availability

All data needed to evaluate the conclusions in the paper are present in the paper and/or the supplementary materials. The dataset generated in this study are available on the following GitHub repository AversiveMemRetrieval. The repository contains all data required to reproduce the results. The raw data are available from the authors upon reasonable request. Source data are provided with this paper.

## Code availability

Codes are available in the AversiveMemRetrieval repository, https://github.com/TheStrangeLab/AversiveMemRetrieval.

## Notes

### Competing Interest Statement

The authors have declared no competing interest.

