## Supplementary Material for "Retrieval of human aversive memories involves reactivation of gamma activity patterns in the hippocampus that originate in the amygdala during encoding"

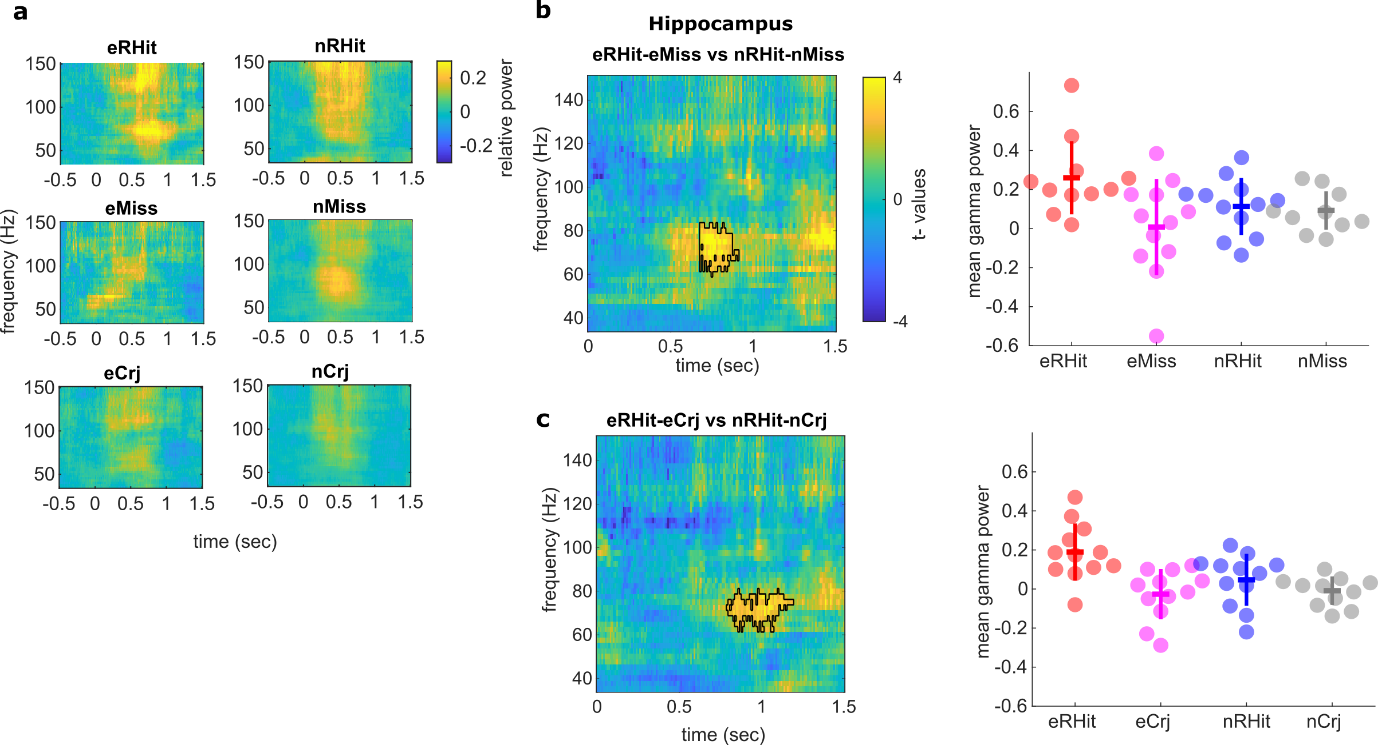
**Supplementary Fig. 1**. **Hippocampal responses during recognition of aversive (e) and neutral (n) scenes.** **a** Time frequency plots showing gamma (35–150 Hz) responses in the hippocampus for aversive and neutral scenes that were remembered (eRHit, nRHit), missed (eMiss, nMiss) and correctly rejected (eCrj, nCrj). **b** Results of time frequency resolved two-sided paired t test . Similarly to the effect shown in the main manuscript (Fig. 1 c), the black outline denotes a first cluster comparing (eRHIt – eMiss and nRHit – nMiss) starting at 0.68s and lasting until 0.91s in the gamma range from 60 to 83Hz, (summed t-value=475.04, P=0.043). The scatter plot indicates the mean hippocampus gamma power values for each subject within the outlined area, as compared to baseline for each condition included in the statistical test **c** The comparison (eRHit – eCrj vs nRHit – nCrj) shows a first cluster (summed t-value=441.59, P=0.069) from 0.79 s to 1.19 s in a similar gamma range (63-80Hz). The scatter plot indicates the same as above, but for statistical test shown in **c**. Note that both effects shown in **b** and **c** do not survive cluster-based correction (at a two-sided threshold p<0.025).

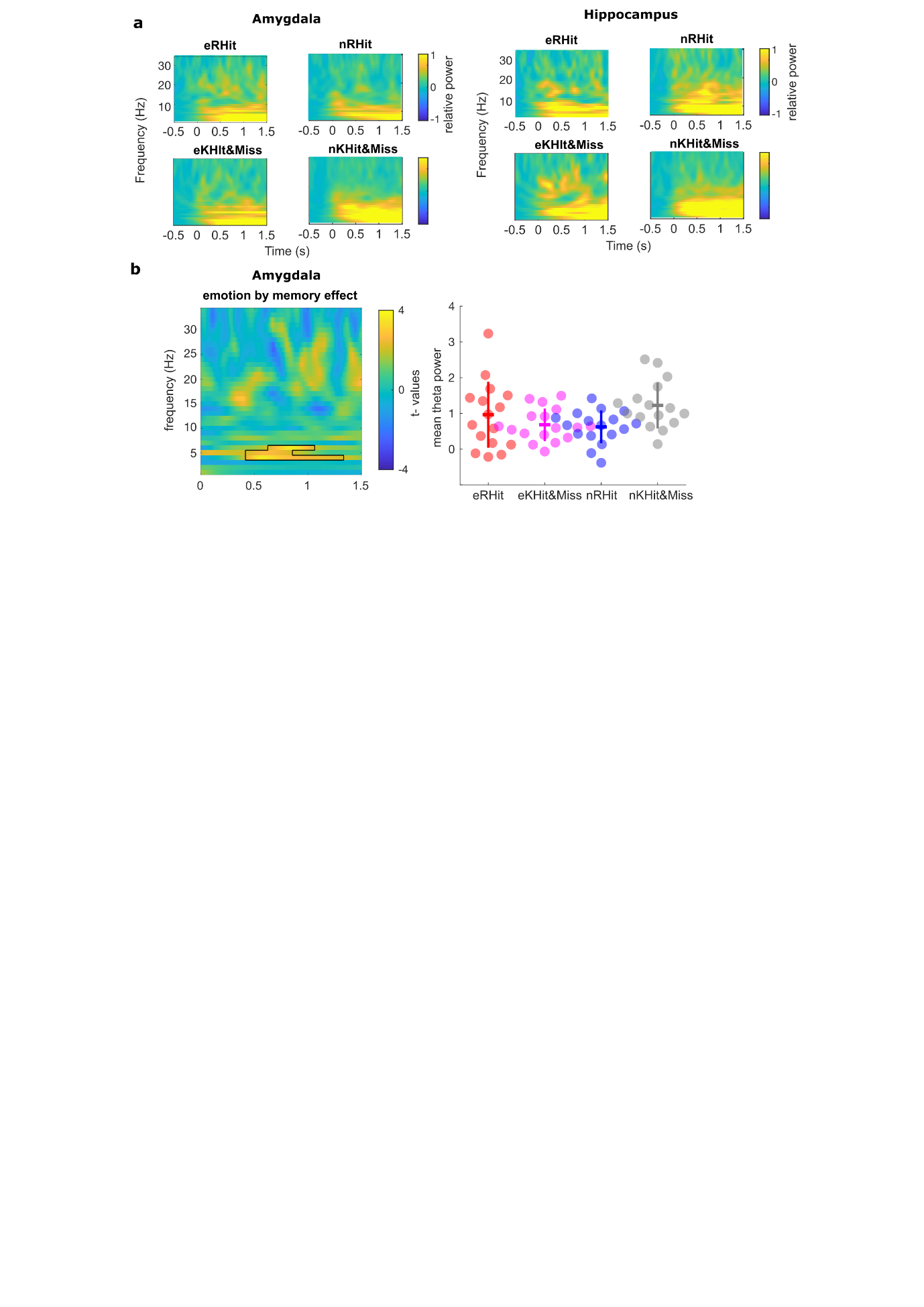

**Supplementary Fig. 2**. **Hippocampal and amygdala low frequency oscillatory power during retrieval of aversive and neutral scenes**. **a** Time frequency plots showing low frequency (0 –34 Hz) responses in the amygdala and the hippocampus for aversive (e ) and neutral (n) scenes remembered (Hit) and correct “know” and incorrect “new” responses (KHit&Miss). **b** Low frequency time-frequency plots showing an amygdala emotion by memory interaction in the theta range (4 - 6 Hz, from 0.4 until 1.3 s), cluster summed t-value=493.11, P=0.016. Post-hoc t-tests on mean power changes across significant time-frequency clusters showed no difference for aversive scene memory (eRHit vs. eKHit&eMiss t_16_ = 1.07, P = 0.29, d = 0.26) but a significant difference for neutral scenes (nRHit vs. nKHit&nMiss t_16_= -3.81, P = 0.0015, d = -0.92). The scatter plot shows the mean theta power values for each subject within the outlined area, as compared to baseline for the four conditions.

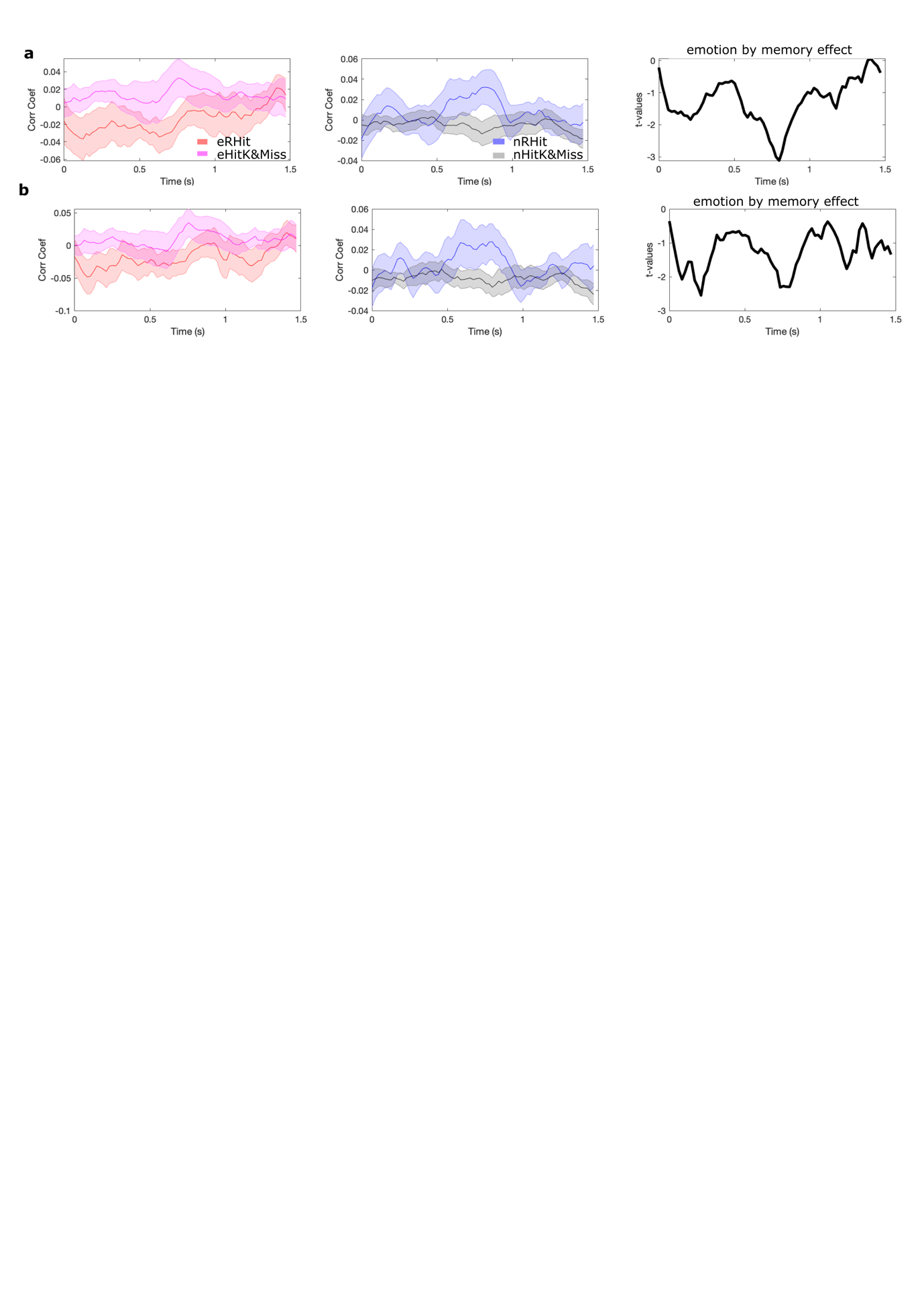
**Supplementary Fig. 3**. **a Amygdala-amygdala Encoding Retrieval similarity (ERS) using representational patterns locked to amygdala high gamma (90-150 Hz) peaks. a** ERS selecting all peaks in each trial and minimum interpeak distance of 0.1s. **b** ERS selecting all peaks in a trial and minimum interpeak distance of 0.3s. No significant emotion by memory ineraction was observed.

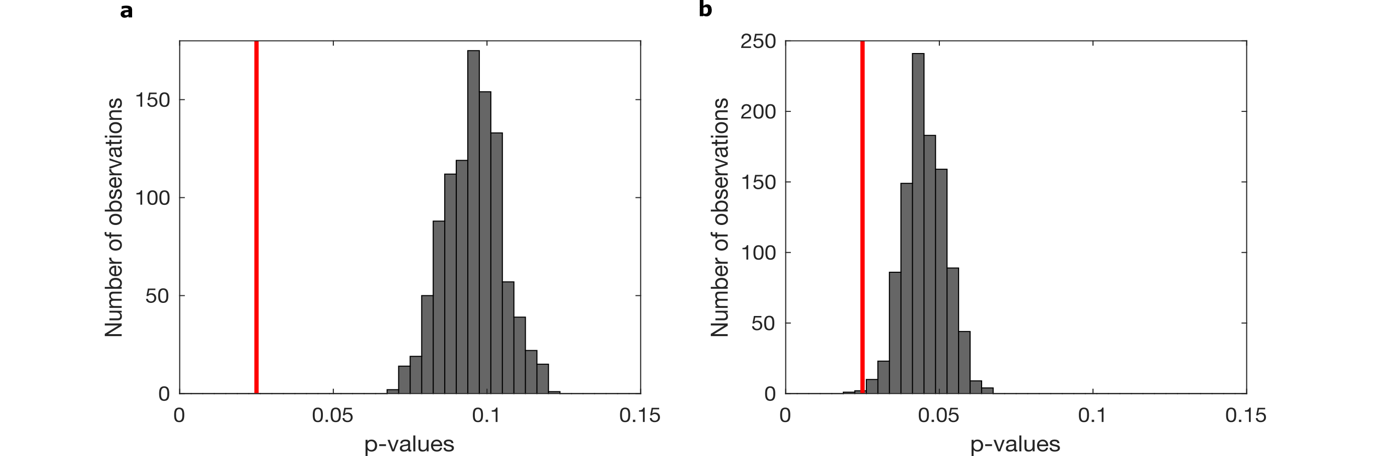
**Supplementary Fig. 4**. **Amygdala-hippocampus EES and ERS random non-peak selection analysis.** We selected as many data segments from the signal as there were peaks in a single trial. These selected segments were of the same time window as used for the peak-related analyses (+/- 0.05 seconds) and were positioned at least 0.03 seconds away from the peak, both before and after it. **a** EES p-values obtained for the emotion by memory interaction using 1000 random non-peak selections. **b** same as in **a** but for the ERS analysis. The red line represent the significant threshold at p=0.025. Both analyses revealed non-significant results. Considered together with the results reported in the main manuscript (Fig. 4), these control analyses highlight the specific relevance of amygdala gamma peaks for hippocampal emotional memory reactivation.

aaa

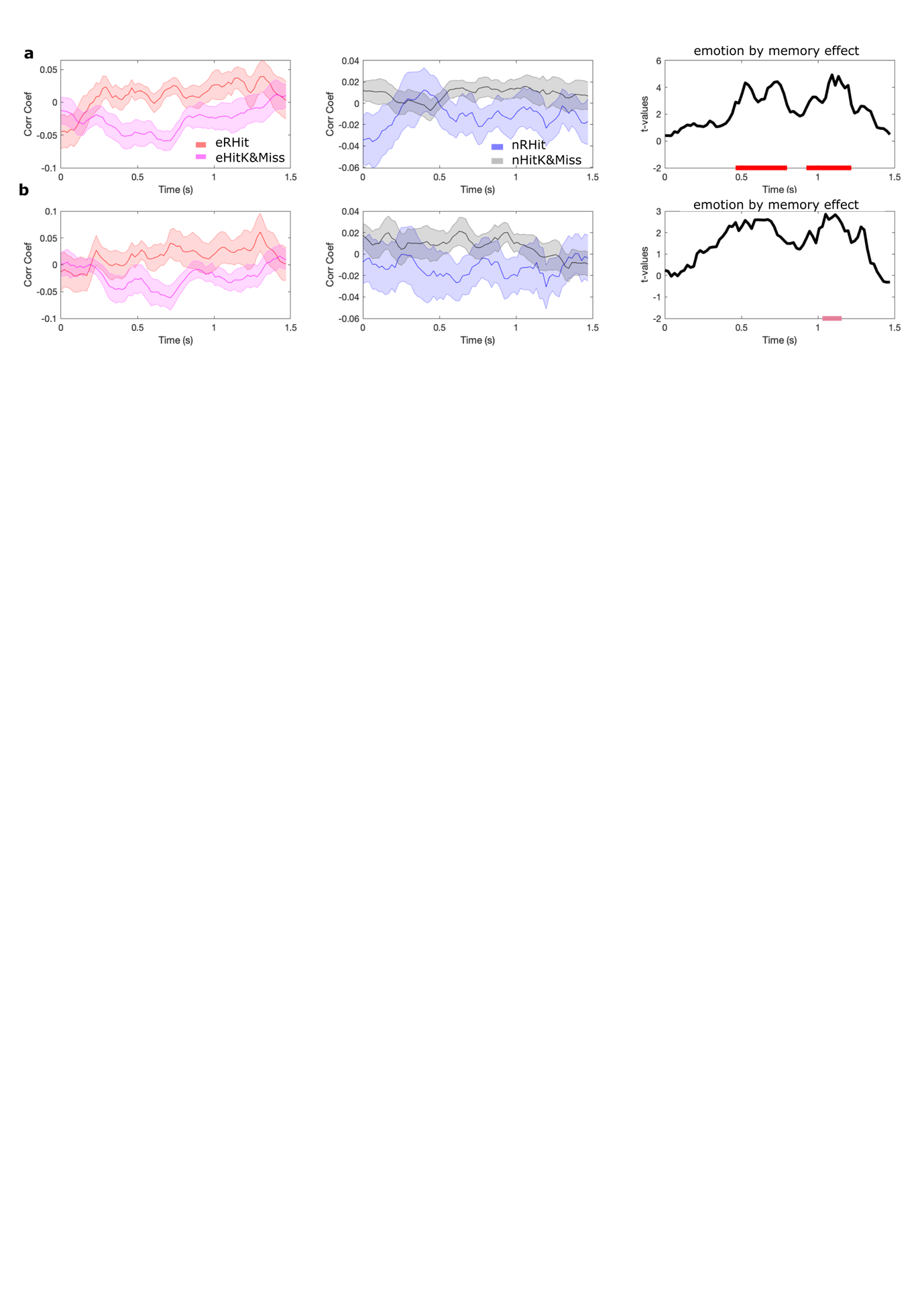

**Supplementary Fig. 5**. **a Amygdala-hippocampus encoding retrieval similarity (ERS) using representational patterns locked to amygdala high gamma (90-150 Hz) peaks. a** ERS selecting all peaks in a trial and minimum interpeak distance of 0.3s. **b** ERS randomly selecting one peak in each trial, and interpeak distance of 0.1s.

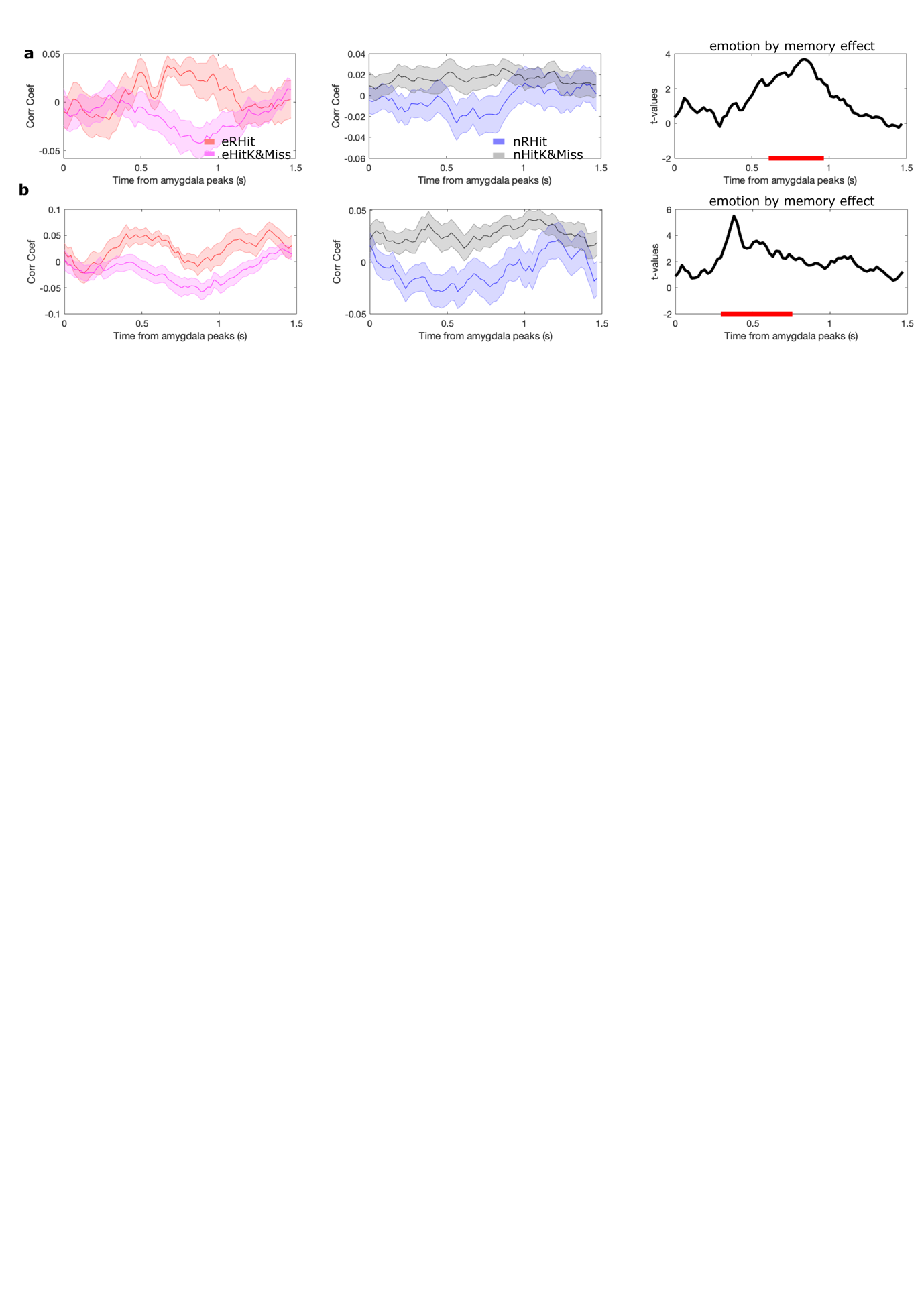

**Supplementary Fig. 6**. **Amygdala-hippocampus encoding encoding similarity (EES) using representational patterns locked to amygdala high gamma (90-150 Hz) peaks. a** EES selecting all peaks in a trial and minimum interpeak distance of 0.3s. **b** EES randomly selecting one peak in each trial, interpeak distance of 0.1s.
